# Acute exposure to caffeine improves navigation in an invasive ant

**DOI:** 10.1101/2023.10.10.561519

**Authors:** Henrique Galante, Massimo De Agrò, Alexandra Koch, Stefanie Kau, Tomer J. Czaczkes

**Affiliations:** Animal Comparative Economics Laboratory, Department of Zoology and Evolutionary Biology, University of Regensburg, 93053 Regensburg, Germany; BRAIR Lab, The BioRobotics Institute, Sant’Anna School of Advanced Studies, 5602 5 Pisa, Italy; Regensburg Center for Biochemistry (RCB), Laboratory for RNA Biology, University of Regensburg, 93053 Regensburg, Germany

**Keywords:** invasion biology, ants, caffeine, navigation, learning, memory

## Abstract

Invasive alien species are a major and growing problem, devastating ecosystems and costing billions of euros in damage and control efforts. Argentine ants, *Linepithema humile*, are particularly concerning, with control efforts often falling short likely due to a lack of sufficient bait consumption. Using neuroactives to manipulate ant navigation and learning could increase recruitment and consumption, ultimately leading to more efficient control strategies. Caffeine is naturally occurring, cheap, and has been found to cause motivational and cognitive improvements in bees. Here, we subject *L. humile* to a wide range of caffeine concentrations and a complex but ecologically relevant task: an open landscape foraging experiment. Without caffeine, we find no effect of consecutive foraging visits on the time the ants take to reach a reward, suggesting a failure to learn the reward’s location. However, low (25ppm) to intermediate (250ppm) concentrations of caffeine lead to a decrease of up to 38% in the time taken to find the reward during each consecutive visit, implying that caffeine boosts learning. Interestingly, such improvements are lost at high (2000ppm) doses. In contrast, caffeine appears to have no impact on the ants’ homing behaviour, as the time required to reach the nest was similar across treatments. The effect of caffeine is thus not only dose-dependent, but also differentially targets neurologically distinct navigational mechanisms. Adding moderate levels of caffeine to baits could be a simple way to improve ant’s ability to learn its location, potentially leading to increased recruitment to, and consumption of, the toxicant.

## Introduction

In Europe, the costs associated with invasive alien species are increasing ten-fold every decade, amounting to billions of euros (Haubrock *et al*., 2021). Among these invasive species, *Linepithema humile* (Mayr, 1868) is particularly damaging, considered one of the most harmful invasive species worldwide (Lowe *et al*., 2000), and the fourth most costly invasive ant species worldwide (Angulo *et al*., 2022). Its impact is not only economic, but also ecological, as they are known to disrupt ecosystems (Silverman & Brightwell, 2008): invasive ants can outcompete native ants, leading to declines in biodiversity and abundance, which in turn, can have a variety of direct and indirect effects on other non-ant species (Holway *et al*., 2002; Alvarez-Blanco *et al*., 2021). Given the severity of their impact, invasive ants are frequently considered a top priority for conservation and pest management programs (Hoffmann *et al*., 2016; Merrill *et al*., 2018; Buczkowski & Wossler, 2019; Suiter *et al*., 2021). Nevertheless, control efforts often fall short, likely due to a lack of sustained bait consumption, suspected to result from competition with natural food sources (Rust, Reierson & Klotz, 2003; Nyamukondiwa & Addison, 2011; Buczkowski, 2016; Lach, Volp & Wilder, 2019).

*L. humile* regularly walk up to twenty thousand times their body length on a foraging trip (Vega & Rust, 2003; Hogg *et al*., 2018). As central place foragers, ants must actively seek out food sources and bring them back to the nest. They do so by employing a highly effective navigation system that minimises their time outside the nest (Collett, Chittka & Collett, 2013; Knaden & Graham, 2016; Buehlmann, Mangan & Graham, 2020). This is likely to increase food acquisition rates (Fewell, 1988; Popp & Dornhaus, 2023) whilst decreasing foraging related risks such as predation and desiccation (Wehner, Marsh & Wehner, 1992; Wehner, Meier & Zollikofer, 2004; Schilman, Lighton & Holway, 2005; Narendra, Reid & Hemmi, 2010). Ant navigation relies predominantly on two mechanisms: path integration and the use of learnt information.

Path integration combines compass information (Wehner & Müller, 2006; Müller & Wehner, 2007; Wajnberg *et al*., 2010; Wystrach *et al*., 2014) with an odometer (Wittlinger, Wehner & Wolf, 2006) to continuously track the ant’s position relative to a reference point, usually either the nest or a frequently visited feeding site (Wehner, 2008). This global vector allows ants to return back to the reference point even when navigating through featureless and novel environments. However, as the distance travelled increases, path integration becomes prone to cumulative errors (Heinze, Narendra & Cheung, 2018). Thus, the path integrator often leads the ant not to an exact point, but to its general vicinity. Once there, they employ well-described systematic searches using spiral search strategies (Müller & Wehner, 1994, 2010; Schultheiss *et al*., 2012; Schwarz, Wystrach & Cheng, 2017) and following CO_2_ plumes (Buehlmann, Hansson & Knaden, 2012) in order to pinpoint the exact location of their nest.

Path integration does not require previous knowledge of the environment and is therefore well suited to naive foragers. However, as ants explore their surroundings, they begin to make use of visual, chemical, and olfactory cues for navigation (Collett *et al*., 1992; Collett, 1996; Aron, Deneubourg & Pasteels, 1988; Czaczkes *et al*., 2014; Arenas & Roces, 2018). Ants learn food-associated cues very rapidly (Roces, 1990; Josens, Eschbach & Giurfa, 2009; Oberhauser *et al*., 2019; Piqueret *et al*., 2022), and are able to integrate multimodal cues (De Agrò *et al*., 2020) and assess their usefulness, prioritising those specific to the nest and not found elsewhere along their route (Huber & Knaden, 2017). In particular, *L. humile* require as little as one experience to successfully associate a location or a scent with a reward (Rossi *et al*., 2020; Wagner *et al*., 2023; Galante & Czaczkes, 2023).

Especially well studied, however, is view-based navigation. Ants learn the visual surroundings of key points by performing specialised learning walks (Collett & Zeil, 2018; Zeil & Fleischmann, 2019), with navigation often improving as the number and range of these increase (Fleischmann *et al*., 2016; Deeti & Cheng, 2021). Furthermore, visual information is continuously gathered along their path to and from food sources. Ants are able to take a panoramic snapshot of a goal, such as their nest, a food source, or a point along a path and later compare it with their current view (Collett & Cartwright, 1983; Cartwright & Collett, 1987; Wystrach & Beugnon, 2009; Wystrach, Beugnon & Cheng, 2011). A misalignment between the memorised panoramic snapshot and the current view leads to gradually increasing image differences. Ants, sensitive to such differences, are able to locate the original snapshot orientation through gradient descent, matching the images by minimizing alignment differences (Collett, Chittka & Collett, 2013). Importantly, view-based navigation differs from path integration in that it does not exclusively rely on idiothetic cues. Instead, it is grounded in an individual’s ability to perceive and learn multimodal cues (Knaden & Graham, 2016). Nevertheless, both mechanisms greatly depend on an individual’s memory retrieval capabilities (Wystrach *et al*., 2020).

Pharmacological interventions offer an excellent tool for dissecting the neural and cognitive basis of behaviours such as navigation (Scheiner *et al*., 2002; Van Swinderen & Andretic, 2011; Felsenberg *et al*., 2011). Moreover, they provide the opportunity to manipulate, and potentially steer, the behaviour of animals. This seems to have already been adopted by various plants, which spike their nectar with secondary metabolites, some of which influence neural activity (Mustard, 2020; Muth *et al*., 2022; Nicolson, 2022). Such chemicals have the potential to artificially manipulate insect behaviour (Baracchi *et al*., 2017; Wink, 2018; Carlesso *et al*., 2021). In this way, deploying neuroactives in artificial baits may be a promising way of enhancing control efficacy.

Mass-recruiting ants, like *L. humile*, rely heavily on pheromone trails to guide nestmates to points of interest (Reid, Latty & Beekman, 2012; Yates & Nonacs, 2016). However, initiating the trails requires pioneering ants to rely on other environmental cues (Knaden & Graham, 2016; Rössler, 2023). Moreover, foraging ants must choose between available pheromone trails, choosing between trails in a manner proportional to their relative strengths (Von Thienen *et al*., 2014). This often results in positive feedback, amplifying initially small differences to result in a collective choice of one path or option (Sumpter & Beekman, 2003; Reid, Sumpter & Beekman, 2011). If a bait additive could enhance path integration, pioneer ants ingesting it might return to their nest faster, likely leading to increased recruitment to the bait relative to scouts which collected food without such additives. This in turn may result in baits with such manipulative additives outcompeting other food sources. Additionally, were the bait additive to improve the acquisition and use of learnt information during navigation, this could result in faster round trips to the bait, with each trip further reinforcing the pheromone trail, increasing recruitment to and consumption of the bait (Czaczkes, Grüter & Ratnieks, 2013).

Caffeine is especially interesting in this regard, as it is naturally occurring, cheap, and well-studied. In honeybees, it has been shown to elicit feeding preference (Singaravelan *et al*., 2005) and to enhance motivation and cognitive performance during complex learning tasks (Si, Zhang & Maleszka, 2005). Additionally, it increased foraging frequency and waggle dancing, quadrupling colony-level recruitment (Couvillon *et al*., 2015). Interestingly, it seems to hinder learning but not memory formation (Mustard *et al*., 2012), although it is reported to improve long-term olfactory associations in honeybees (Wright *et al*., 2013). Similarly, caffeine improved bumblebee odour association learning (Arnold *et al*., 2021) and increased flower pollination (Thomson, Draguleasa & Tan, 2015). However, it lowered bumblebee food consumption (Tiedeken *et al*., 2014). In ants, it likely alters food value perception, acting as both an attractant and a repellent, depending on the plant extract and concentration used (Yeoh, Dieng & Majid, 2018; Majid *et al*., 2018; Madsen & Offenberg, 2019). Furthermore, caffeine has been reported to improve learning and memory in *Myrmica sabuleti* ants, albeit at the cost of decreased food consumption (Cammaerts, Rachidi & Gosset, 2014).

Contrastingly, recent work suggests a general lack of effect of an intermediate concentration of caffeine on both spatial and olfactory associative learning in *L. humile* (Galante & Czaczkes, 2023). However, that study focused on a single concentration of caffeine and overlooked its potential effects on navigation. Additionally, it tested learning in a binary choice Y-maze setup, which proved to be a very easy task for the ants, leading to a ceiling effect which further obscured the potential effects of caffeine. Here, we subject *L. humile* to a wider range of caffeine concentrations and a more complex and ecologically relevant task: an open landscape foraging experiment. By using an automated tracking system, we collected high resolution data of different segments of the foraging journey, as opposed to simple associative learning binary choices, studying not only the effect of caffeine on learning and memory, but also on locomotion and navigation.

## Materials and Methods

### Colony maintenance

*Linepithema humile* (Mayr, 1868) were collected from Portugal (Proença-a-Nova) and Spain (Girona) between April 2021 and January 2022. Ants were split into colony fragments (henceforth colonies), containing three or more queens and 200-1000 workers, kept in non-airtight plastic boxes (32.5 x 22.2 x 11.4 cm) with a plaster of Paris floor and PTFE coated walls. 15mL red transparent plastic tubes, partly filled with water, plugged with cotton, were provided as nests. Ants were maintained on a 12:12 light:dark cycle at room temperature (21-26 °C) with *ad libitum* access to water. Between experiments, ants were fed *ad libitum* 0.5M sucrose solution and *Drosophila melanogaster* twice a week. During experiments, ants were fed once a week and deprived of carbohydrates for four to five days prior to testing, ensuring high foraging motivation. Experiments were conducted between September 2021 and March 2022 using 14 colonies divided into donor/recipient pairs. Focal ants left their original colony (donor) but returned to a different colony (recipient) to unload the contents of their crop. This ensured donor colonies, and consequently focal ants, were never exposed to caffeine prior to the experiment.

### Chemicals and solutions

Caffeine (CAS 58-08-2) was obtained from Sigma-Aldrich (Taufkirchen, Germany). 1M sucrose solutions (Südzucker AG, Mannheim, Germany) mixed with different caffeine concentrations were used as treatments. Identical 1M sucrose solutions were used as controls. Caffeine concentrations were chosen based on previous reports of their effects on Hymenopterans. Caffeine solutions ranging from naturally occurring concentrations (Singaravelan *et al*., 2005; Couvillon *et al*., 2015) to those reported as the LD_50_ of honeybees (Detzel & Wink, 1993) were used: 25ppm (0.13μmol ml^-1^), 250ppm (1.29μmol ml^-1^) and 2000ppm (10.30μmol ml^-1^), respectively. A double-blind procedure was applied to all solutions used in order to minimize experimenter bias.

### Open landscape spatial learning

A challenging and ecologically relevant experimental paradigm was developed to study the effects of different concentrations of caffeine on spatial learning and memory. This setup consisted of an A4 (210 x 297 mm) disposable paper overlay platform, surrounded by a water moat, with a single 1cm entrance, attached to a fixed top-view Raspberry Pi HQ camera setup. A sucrose solution drop (positive stimulus), either pure or laced with 25ppm, 250ppm or 2000ppm of caffeine, was positioned at the entrance of the platform (see blue drop in Figure 1B). A single ant from a donor colony was allowed onto the platform via a drawbridge, was marked with acrylic paint while drinking, and returned to its paired recipient colony. Meanwhile, the disposable overlay was replaced, ensuring the removal of any pheromone trails laid, and a drop of the solution used during the first visit placed on one of the sides of the platform (see green drops in Figure 1B). After unloading, the ant was placed back onto the platform’s entrance where it was video recorded and the time it spent in the colony, the time it took to reach the reward, to drink and to return to the colony was noted for each visit. Overall, each ant carried out five consecutive visits to the platform: a priming visit, with the reward placed at the entrance of the platform, followed by four visits with the reward either always on the left or the right side of the landscape. 142 individuals were tested, with a complete run taking between 35 to 155 minutes.

**Figure 1.**
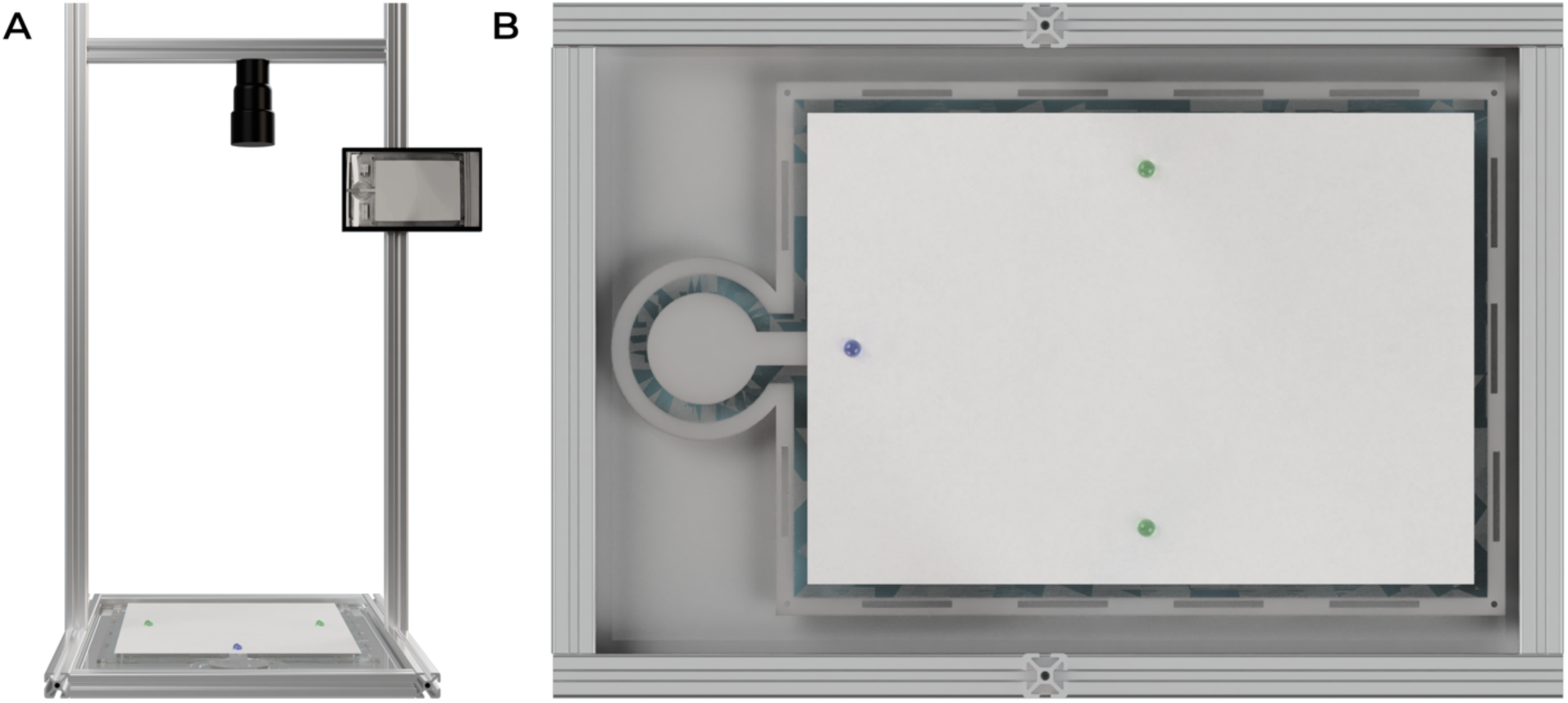
– Open landscape experimental setup. **(A)** Front view. A4 (210 x 297 mm) disposable paper overlay platform, surrounded by a water moat, with a single 1cm entrance, attached to a fixed top view Raspberry Pi HQ camera setup. **(B)** Top view. Blue drop represents the location of the sucrose solution drop (positive stimulus), either the control or the treatment on the ant’s first visit (2cm from the middle of the platform’s entrance). Green drops represent the possible location of the positive stimulus during the remaining visits (2.5cm from either the left or right side of the platform – each ant experienced the stimulus exclusively on one side).

### Data extraction and processing

DeepLabCut version 2.2.2 (Mathis *et al*., 2018; Nath *et al*., 2019) was used to automate video analysis, allowing for the acquisition of coordinates for each ant’s head and gaster, as well as those of each corner of the A4 platform and the centre of the solution drop at every frame (videos recorded at 30fps). In total 142 ants were recorded, however five of them were excluded as they were unable to find the reward within 20 minutes in at least two of their five visits to the landscape. Furthermore, 11 ants failed to locate the reward in one of their five visits and one other was lost in the colony before it could do its final visit, therefore those visits were removed. 10 other visits were excluded due to tracking associated issues (specified in the detailed statistical analysis and code). Overall, 526 visits of 136 ants were analysed. Python version 3.7.13 (Rossum & Drake, 2009) was used to standardise the ants’ coordinates by ensuring the same corner of the A4 platform was used as the origin of the cartesian referential of all videos. The known dimensions of the A4 were further used to convert coordinates from pixels to millimetres. To account for DeepLabCut tracking errors, any ant movement exceeding two millimetres per frame was considered implausible and subsequently removed. This is due to biological constraints that limit ants’ movement to a maximum speed of around one millimetre per frame (Burford *et al*., 2018). In this process, an average of 0.1% [0%, 0.8%, N = 526] of frames were eliminated from each video. The times at which an ant reached and left the reward were automatically derived from the tracking data. These were used to define the ant’s foodward and nestward journeys. The foodward journey encompassed the time from when the ant initially entered the landscape until it reached the reward for the first time. The nestward journey was taken as the last time the ant left the reward until it left the landscape. For each visit, the mean instantaneous speed and path tortuosity were then calculated separately for both the foodward and nestward journeys. Mean instantaneous speed represents the average distance travelled by the ant within a frame (1/30s), while path tortuosity quantifies the total distance covered by the ant relative to the minimum possible distance it could have travelled, providing a measure of path straightness. Importantly, exclusion criteria and extracted variables were defined *a priori* in a pre-registration with a few notable deviations. Mainly, the 25ppm and 250ppm caffeine treatments were added at a later stage and the second visit to the landscape was analysed together with all others in order to better understand the effect of consecutive visits on navigation.

### Statistical analysis

The complete statistical analysis output, and the entire dataset on which this analysis is based, is available from Zenodo (https://doi.org/10.5281/zenodo.8413980).

All graphics and statistical analysis were generated using R version 4.2.1 (Wickham, 2016; Kassambara, Kosinski & Biecek, 2021; R Core Team, 2022; Wickham, 2022). Foodward and nestward journey duration were analysed using mixed effects cox proportional-hazards models (Therneau & Grambsch, 2000; Therneau, 2022). Note that for the survival analysis of foodward journey duration the 11 ants which failed to find the reward in one of their five visits were included yet censored. Mean instantaneous speed and path tortuosity were analysed with linear mixed-effects models (Bates *et al*., 2015). DHARMa (Hartig, 2022) was used to assess linear model assumptions and MuMIn (Bartoń, 2022) to obtain a measure of goodness of fit. Analysis of variance tables were used to test the effects of the regressions coefficients (Fox & Weisberg, 2019). Estimated marginal means of linear trends and contrasts were obtained using the emmeans package (Lenth, 2022) with Bonferroni adjusted values accounting for multiple testing. Consecutive visit effects were assumed to be relatively linear as previously shown in Y-maze experiments performed on *L. humile* (Galante & Czaczkes, 2023; Wagner *et al*., 2023). We avoid the use of p-values, and their associated binary decision of significant/nonsignificant, instead reporting effect size estimates and their respective 95% confidence intervals shown throughout the results section as (estimate [lower limit, upper limit, N = sample size]).

## Results

There was no clear effect of caffeine concentration and/or the number of consecutive foraging visits to the landscape on the time ants spent drinking the reward (84s [min = 71s, max = 107s]).

### Low to intermediate caffeine concentrations lead to foodward path optimisation

The time it took for an ant to reach the reward upon entering the landscape was influenced by both caffeine concentration and the number of consecutive visits to the landscape, as well as the interaction between the two (Figure 2). On average, across treatments and visits, the foodward journey took 212s [min = 63s, max = 372s].

**Figure 2.**
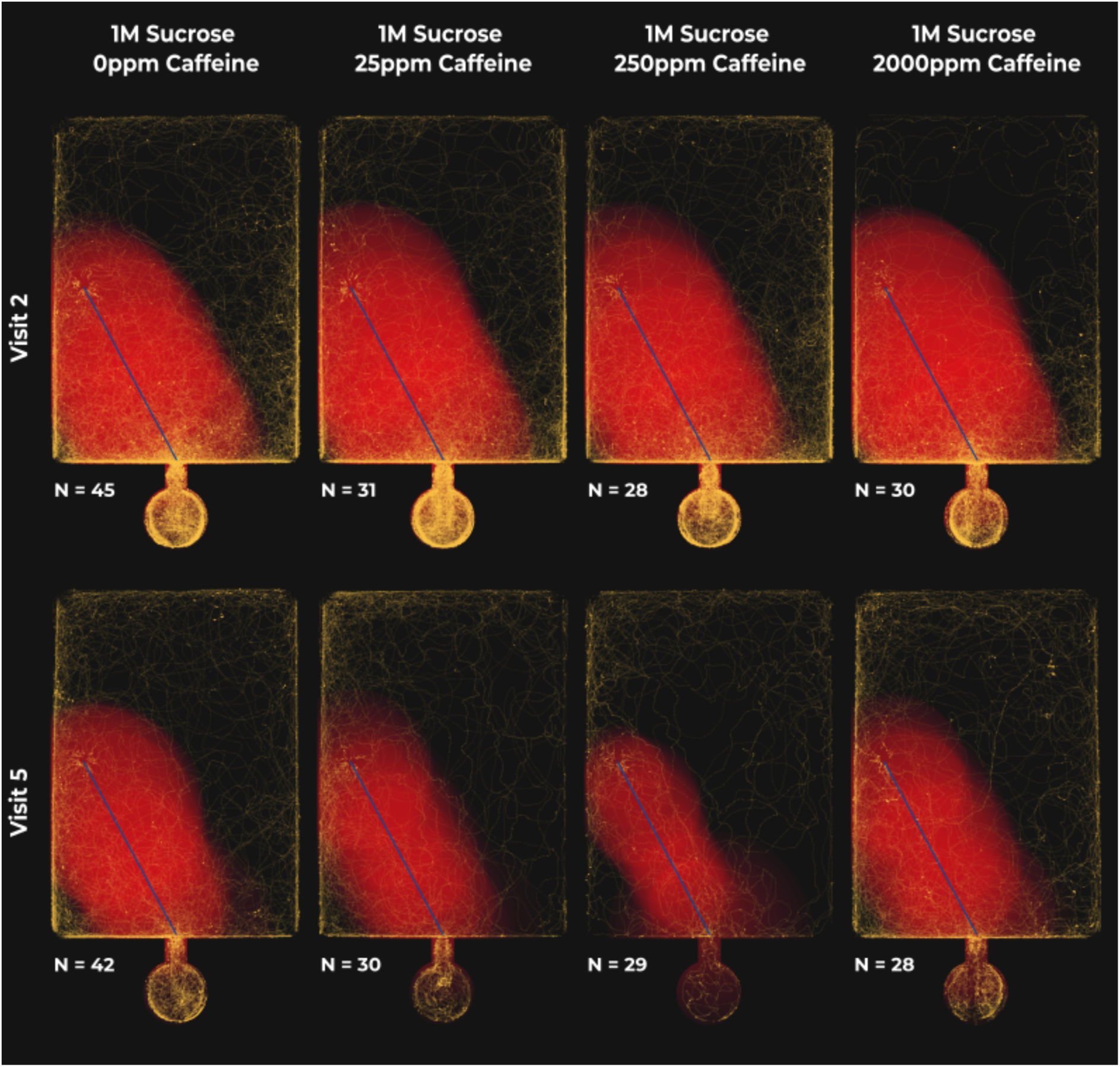
– Dose-dependent effect of caffeine on consecutive foodward visits to the open landscape. Red areas highlight the average deviation of all ant paths from the shortest route (blue line) at each time point (1% increments = 12s increments at most). Time normalisation sets the starting point of each path at 0% and the final point at 100%. Larger areas surrounding the shortest path indicate larger deviations from it, while brighter regions suggest greater overlap between deviations at different time points implying a lack of directional movement. Note that closer adherence to the straightest line results in smaller, dimmer red areas, which translates into shorter times spent in the landscape and therefore fewer yellow points on the graph. Yellow points highlight the paths taken by individual ants, from first entering the landscape until first reaching the reward, with brighter spots indicating overlapping trajectories. To aid visualization, the paths of ants reaching the reward on the right side of the landscape were mirrored.

Under control conditions, ants are likely to be 5.6% [-12.6%, 23.8%, N = 537] faster at reaching the reward with every consecutive visit. Meaning, if an ant initially took 300s to find the reward, after three consecutive visits to the open landscape, used henceforth as the example, it is likely to find the reward in 252s [133s, 428s]. However, when exposed to 25ppm of caffeine, consecutive visits result in a 27.8% [5.7%, 50.0%, N = 537] likelihood of ants having a faster foodward journey (113s [38s, 252s]). This effect is almost doubled in the 250ppm caffeine treatment, where ants are likely to find the reward 43.5% [19.5%, 67.6%, N = 537] faster with every consecutive visit (54s [10s, 156s]). However, the benefits observed at lower to intermediate caffeine concentrations are lost when ants are fed 2000ppm of caffeine (3.1% [– 20.5%, 26.8%, N = 537] | 273s [118s, 525s]). Furthermore, the effect of consecutive visits is 38.0% [7.8%, 68.2%, N = 537] higher in ants under the 250ppm caffeine treatment when compared to control-treated ants and 40.4% [6.6%, 74.2%, N = 537] higher when compared to the 2000ppm caffeine treatment (Figure 3).

**Figure 3.**
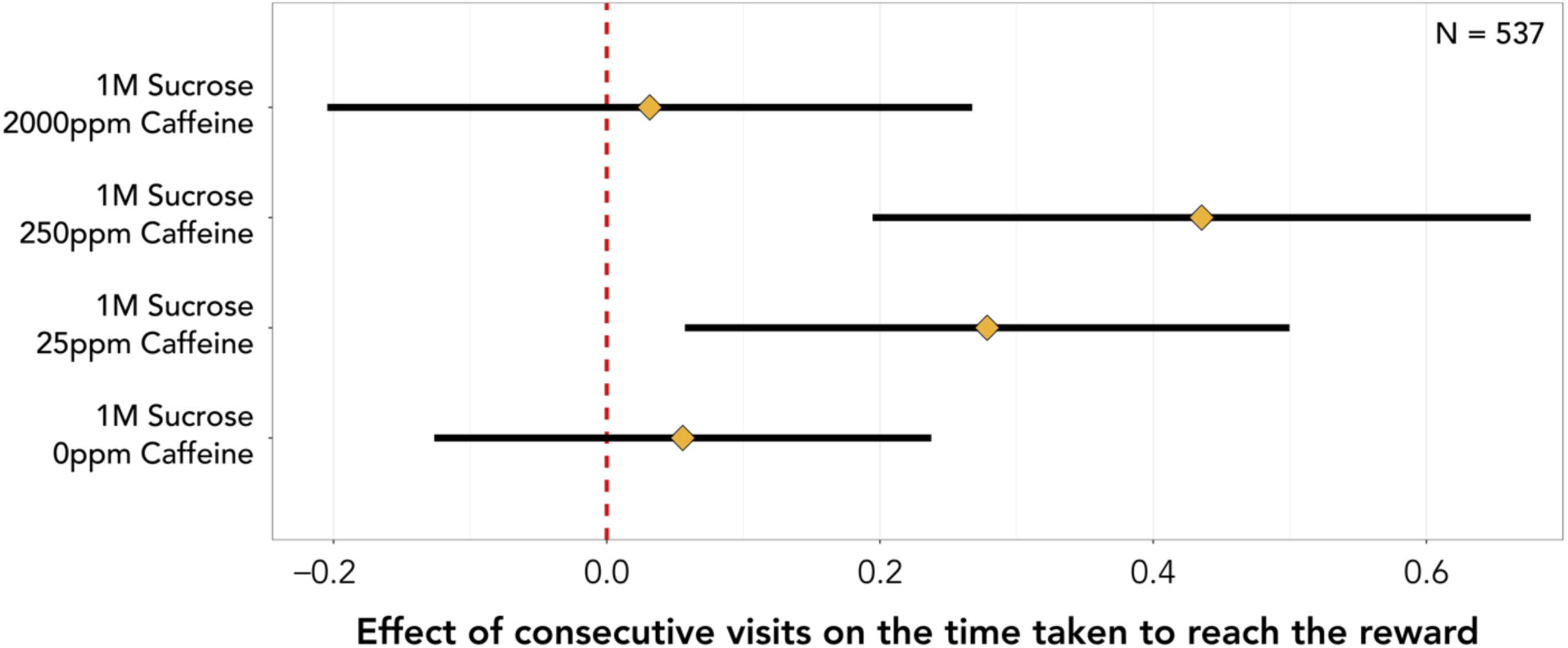
– Effect of consecutive visits on the time an ant takes to reach the reward since first entering the landscape for each caffeine treatment. Diamonds represent the estimated marginal means of linear trends obtained from the mixed effects cox proportional-hazards model and whiskers the respective 95% confidence intervals. Estimates of 0 (red dashed vertical line) indicate no effect of consecutive visits, whilst estimates > 0 or < 0 indicate that ants are more or less likely, respectively, to reach the reward faster with consecutive visits. If the 95% confidence intervals include an estimate of 0 it is likely that there is no effect of consecutive visits.

Notably, caffeine did not affect mean instantaneous speed during the foodward journey. However, from the first (17.5 mm/s [min = 9.0 mm/s, max = 25.5 mm/s, N = 134]) to the last (18.6 mm/s [min = 7.4 mm/s, max = 27.7 mm/s, N = 129]) visit to the landscape, mean instantaneous speed increased by 0.3 mm/s [0.1 mm/s, 0.6 mm/s, N = 526] per visit. The logarithm of path tortuosity decreased by 0.4 [0.0, 0.8, N = 526] with each consecutive visit under control conditions and by 0.4 [-0.1, 0.9, N = 526] under 2000ppm caffeine treatment. 25ppm (0.9 [0.4, 1.4, N = 526]) and 250ppm caffeine treatments (0.8 [0.3, 1.3, N = 526]) exhibited a similar, yet stronger effect. Importantly, effects in the logarithmic scale are often exponential in the linear scale. Meaning, if an ant had a path tortuosity of 20 on its first visit to the landscape, after three consecutive visits, we would expect it to have a path tortuosity of 6.0 [1.8, 20.0] under the control treatment, and a path tortuosity of 1.8 [1.0, 8.1] under the 250ppm caffeine treatment.

### Caffeine had no effect on the nestward journey

The time it took for an ant to exit the landscape upon last touching the reward was influenced by the number of consecutive visits to the landscape, but not by the amount of ingested caffeine (Figure 4). On average, across treatments and visits, the nestward journey took 44s [min = 30s, max = 57s].

**Figure 4.**
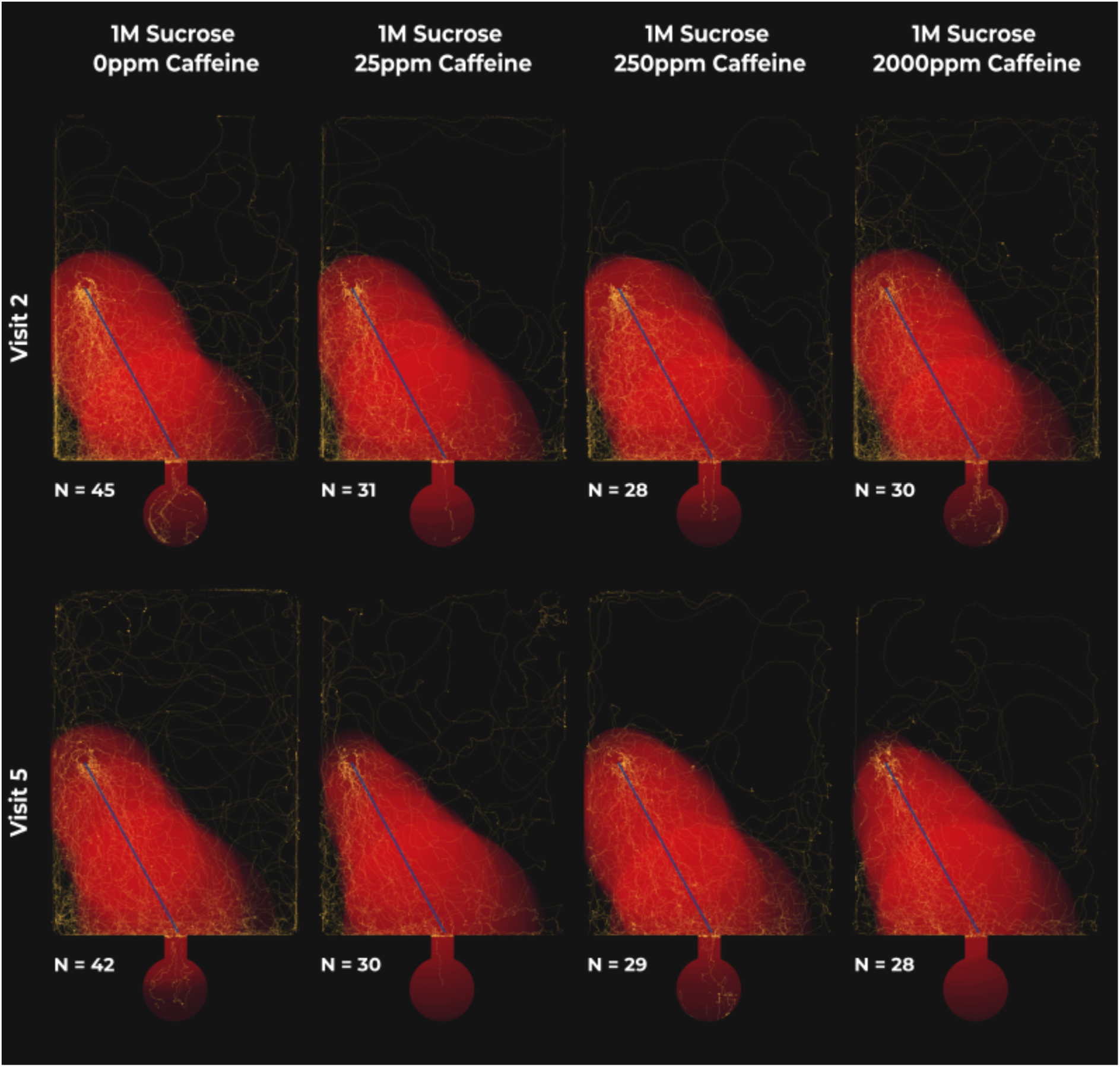
– No effect of caffeine on consecutive nestward journeys. Red areas highlight the average deviation of all ant paths from the shortest route (blue line) at each time point (1% increments = 2s increments at most). Time normalisation sets the starting point of each path at 0% and the final point at 100%. Larger areas surrounding the shortest path indicate larger deviations from it, while brighter regions suggest greater overlap between deviations at different time points implying a lack of directional movement. Note that closer adherence to the straightest line results in smaller, dimmer red areas, which translates into shorter times spent in the landscape and therefore fewer yellow points on the graph. Yellow points highlight the paths taken by individual ants, from last touching the reward until leaving the landscape, with brighter spots indicating overlapping trajectories. To aid visualization, the paths of ants presented a reward on the right side of the landscape were mirrored.

With each consecutive visit ants are likely to be 11.0% [2.8%, 19.2%, N = 526] faster at returning to the nest. Meaning, if an ant initially took 30s to return to its nest, after three consecutive visits to the open landscape, it is likely to return in 21s [16s, 28s], regardless of its consumption of caffeine (Figure 5). Contrastingly, mean instantaneous speed (14.5 mm/s [min = 4.9 mm/s, max = 26.9 mm/s, N = 526]) was constant throughout consecutive visits and treatments. Furthermore, albeit unaffected by caffeine, the logarithm of path tortuosity decreased by 0.07 [0.02, 0.11, N = 526] per visit. Therefore, if an ant had a path tortuosity of 4 on its first visit to the landscape, after three consecutive visits, we would expect it to have a path tortuosity of 3.2 [2.9, 3.8].

**Figure 5.**
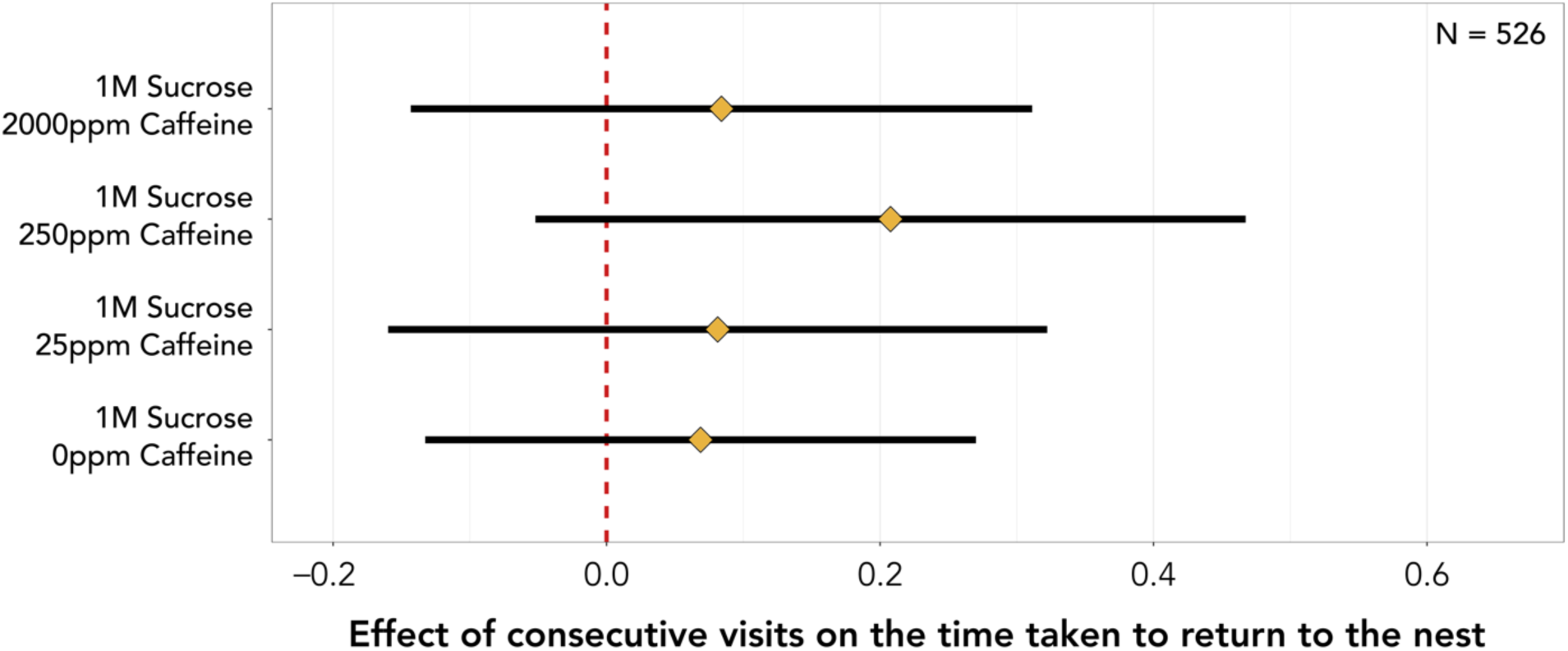
– Effect of consecutive visits on the time an ant took to return to the nest since leaving the reward for the last time for each caffeine treatment. Diamonds represent the estimated marginal means of linear trends obtained from the mixed effects cox proportional-hazards model and whiskers the respective 95% confidence intervals. Estimates of 0 (red dashed vertical line) indicate no effect of consecutive visits, whilst estimates > 0 or < 0 indicate that ants are more or less likely, respectively, to return to the nest faster with consecutive visits. If the 95% confidence intervals include an estimate of 0 it is likely that there is no effect of consecutive visits.

## Discussion

Low (25ppm) and intermediate (250ppm) concentrations of caffeine shortened the foodward journey over consecutive visits (Figure 2 and 3). This was not due to an increase of mean instantaneous speed, but rather an increase in path straightness over consecutive visits under these treatments. Thus, caffeine is likely dose-dependently boosting learning, as a straighter path suggests the ant knows the location of the reward (Figure 2). However, the improvements gained were lost at high doses of caffeine (2000ppm). This suggests a hormetic dose-response pattern, where caffeine is toxic at high doses, but when ingested in smaller amounts has the opposite effect, stimulating biological function. Similar results were previously found in bees for a variety of chemicals (Cutler & Rix, 2015), including caffeine (Wright et al., 2013). In humans, low to moderate caffeine dosages often have stimulant and performance-enhancing effects. On the other hand, high doses may be associated with aversive somatic effects, including sleep disruption and increased anxiety and agitation, all of which can contribute to impaired fine motor control (Kaplan *et al*., 1997; Smith, 2002; Pallarés *et al*., 2013; Souza *et al*., 2022). In honeybees, higher doses of caffeine have resulted in a reduced likelihood of bees to respond to a conditioned odour (Mustard *et al*., 2012) and 2000ppm of caffeine has previously been found as the LD_50_ of honeybees (Detzel & Wink, 1993). Thus, it seems reasonable that the loss of effect at high doses of caffeine is likely due to its toxicity.

Interestingly, caffeine had no clear impact on nestward journey duration (Figures 4 and 5). Much as in the foodward journey, consecutive visits did not alter the speed at which the ants moved. However, ants were, on average, slower during the nestward journey when compared to the foodward journey, possibly due to being heavier when their crop is full. Independently of caffeine, over consecutive visits nestwards paths became straighter, which translated into faster nestward journeys across treatments. The lack of effect of caffeine during the nestward journey, paired with the fact that foodward journey durations were an order of magnitude slower than nestward journey durations, suggests different navigational mechanisms are being selectively used throughout the foraging journey. Path integration, being innate, constantly active, and not requiring learning of the surrounding landscape (Zeil, Narendra & Stürzl, 2014; Zeil, 2022), is likely to be extremely accurate under relatively short distances such as the ones travelled in the open landscape (Heinze, Narendra & Cheung, 2018). It is thus likely to be the main mechanism used during the nestward journey, albeit paired with a learnt component, as nestward journey duration did improve over consecutive visits. In their natural environment, ants walk longer distances while foraging, and face a more complex environment with at times contradicting multimodal cues. Under such conditions, where learning and memory play a key role, caffeine might have a greater impact on the nestward journey. In fact, we note a slight trend towards faster nestward journeys over consecutive visits in the 250ppm treatment (Figure 5).

Consecutive visits did not alter foodward journey duration in control treated ants (Figure 3). This implies that three visits to the reward were not sufficient for the ants to learn and/or memorise its location. However, previous work has demonstrated that *L. humile* are excellent spatial and olfactory learners often requiring as little as one visit to the reward to learn its characteristics and location (Rossi *et al*., 2020; Wagner *et al*., 2023; Galante & Czaczkes, 2023). We were thus successful in creating a more complex and challenging experiment, which arguably better depicts natural conditions. Furthermore, this enabled us to exclude any ceiling effects potentially observed in previous experiments which might have masked any effects resulting from exposure to caffeine (Galante & Czaczkes, 2023).

The range over which panoramic views offer navigational guidance is smaller in denser environments and larger in open landscapes (Zeil, Hofmann & Chahl, 2003). Considering we purposefully removed landmarks from the experiment, ants are likely to have heavily relied on positional image matching during the foodward journey. In their invasive range, and considering the temporary nature of their nests, *L. humile* habitats are likely to be richer in landmarks. Therefore, ants will not only rely on visual cues but also olfactory ones. In fact, ants have been shown to use olfactory landmarks to locate food and their nest (Helmy & Jander, 2003; Steck, Hansson & Knaden, 2009; Huber & Knaden, 2018) with a combination of visual and olfactory navigational tools leading to increased performance (Steck, Hansson & Knaden, 2011). Caffeine has been shown to improve olfactory associations in both honeybees and bumblebees (Wright *et al*., 2013; Arnold *et al*., 2021). It is thus likely that the observed navigational improvements of caffeine will not only be maintained, but potentially increased, in invaded areas as ants need to navigate more complex environments relying on multimodal cues.

Similarly to Si, Zhang & Maleszka, (2005), we found that caffeine has no effect in a simple Y-maze spatial and olfactory association learning task (Galante & Czaczkes, 2023), but its effects become visible in a more complex and field-realistic experiment. Low to intermediate caffeine concentrations are likely to act by enhancing alertness and motivation. This may be due to the effects of caffeine as an adenosine receptor antagonist, potentially leading to increased levels of acetylcholine which might improve cognitive alertness (Gold, 2003). Moreover, a decrease of GABA, a neurotransmitter thought to impair olfactory memory in bees (Boitard *et al*., 2015), might reduce forgetting of the location of the caffeinated rewards. In turn, this could generate addiction-like symptoms potentially increasing the ant’s motivation to forage on such rewards. However, a recent study exploring visual learning in bumblebees shows GABA promoting flower fidelity (Calderai *et al*., 2023). Caffeine has been shown to increase responsiveness and activity in jumping spiders (Humphrey, Helton & Nelson, 2019) as well as having an effect on locomotion in a variety of insects (Mustard, 2014). However, we show no effect of caffeine on ant mean instantaneous speed throughout the foraging journey, suggesting caffeine-mediated navigational improvements are not a consequence of motor efficiency but rather of cognitive enhancement.

Adding neuroactive substances to toxic baits may hold potential as a novel approach to improve invasive insect control. Future work, ideally under natural conditions, should test whether caffeine interacts with the chosen toxicant, and if it also improves recruitment and visitation rates in ants, as previously shown in honeybees and bumblebees, respectively (Couvillon *et al*., 2015; Thomson, Draguleasa & Tan, 2015). As new invasive ant species start to become established worldwide (Menchetti *et al*., 2023), there is growing urgency in finding ways to improve control efforts. Adding low dosages of caffeine to toxic baits may be a cheap and easily deployable method of boosting learning of bait location, potentially leading to increased recruitment to and consumption of the toxicant, and ultimately improved control.

## Acknowledgements

We thank E. Sequeira and S. Abril for ant collection, and A. C. LeBoeuf for suggestions on graphical visualisation.

## Funding

H. Galante and M. De Agrò were supported by an ERC Starting Grant to T. J. Czaczkes (H2020-EU.1.1. #948181). T. J. Czaczkes was supported by a Heisenberg Fellowship from the Deutsche Forschungsgemeinschaft (CZ 237 / 4-1). A. Koch was supported by a University of Regensburg “Anreizsystem” grant to T. J. Czaczkes.

## Declaration of interests

The authors declare no competing interests.

## Ethical statement

We have conducted all experiments in accordance with the guidelines that are applicable to working with the model organism in the European Union. Colonies were kept in closed boxes under oil baths in order to prevent any escape.

## Author contributions: H. Galante

Conceptualization, Methodology, Software, Validation, Formal analysis, Investigation, Data Curation, Writing – original draft, Writing – review & editing, Visualization, Supervision. **M. De Agrò:** Methodology, Validation, Writing – review & editing. **A. Koch:** Investigation, Writing – review & editing. **S. Kau:** Investigation, Writing – review & editing. **T. J. Czaczkes:** Conceptualization, Methodology, Resources, Writing – review & editing, Supervision, Project administration, Funding acquisition.

## Notes

https://doi.org/10.5281/zenodo.8413980

### Competing Interest Statement

The authors have declared no competing interest.

## References

1. Alvarez-Blanco, P., Cerdá, X., Hefetz, A., Boulay, R., et al. (2021) Effects of the Argentine ant venom on terrestrial amphibians. Conservation Biology. 35 (1), 216–226. 10.1111/cobi.13604.

2. Angulo, E., Hoffmann, B.D., Ballesteros-Mejia, L., Taheri, A., et al. (2022) Economic costs of invasive alien ants worldwide. Biological Invasions. 24, 2041–2060. 10.1007/s10530-022-02791-w.

3. Arenas, A. & Roces, F. (2018) Appetitive and aversive learning of plants odors inside different nest compartments by foraging leaf-cutting ants. Journal of Insect Physiology. 109, 85–92. 10.1016/j.jinsphys.2018.07.001.

4. Arnold, S.E.J., Dudenhöffer, J., Fountain, M.T., James, K.L., et al. (2021) Bumble bees show an induced preference for flowers when primed with caffeinated nectar and a target floral odor. Current Biology. 31 (18). 10.1016/j.cub.2021.06.068.

5. Aron, S., Deneubourg, J.L. & Pasteels, J.M. (1988) Visual cues and trail-following idiosyncrasy in *Leptothorax unifasciatus*: An orientation process during foraging. Insectes Sociaux. 35 (4), 355–366. 10.1007/BF02225811.

6. Baracchi, D., Marples, A., Jenkins, A.J., Leitch, A.R., et al. (2017) Nicotine in floral nectar pharmacologically influences bumblebee learning of floral features. Scientific Reports. 7 (1). 10.1038/s41598-017-01980-1.

7. Bartoń, K. (2022) MuMIn: Multi-model inference. Available from: https://CRAN.R-project.org/package=MuMIn.

8. Bates, D., Mächler, M., Bolker, B. & Walker, S. (2015) Fitting linear mixed-effects models using lme4. Journal of Statistical Software. 67 (1), 1–48. 10.18637/jss.v067.i01.

9. Boitard, C., Devaud, J., Isabel, G. & Giurfa, M. (2015) GABAergic feedback signaling into the calyces of the mushroom bodies enables olfactory reversal learning in honey bees. Frontiers in Behavioral Neuroscience. 9. 10.3389/fnbeh.2015.00198.

10. Buczkowski, G. (2016) The Trojan horse approach for managing invasive ants: a study with Asian needle ants, Pachycondyla chinensis. Biological Invasions. 18 (2), 507–515. 10.1007/s10530-015-1023-z.

11. Buczkowski, G. & Wossler, T.C. (2019) Controlling invasive Argentine ants, *Linepithema humile*, in conservation areas using horizontal insecticide transfer. Scientific Reports. 9 (1). 10.1038/s41598-019-56189-1.

12. Buehlmann, C., Hansson, B.S. & Knaden, M. (2012) Path integration controls nest-plume following in desert ants. Current Biology. 22 (7), 645–649. 10.1016/j.cub.2012.02.029.

13. Buehlmann, C., Mangan, M. & Graham, P. (2020) Multimodal interactions in insect navigation. Animal Cognition. 23 (6), 1129–1141. 10.1007/s10071-020-01383-2.

14. Burford, B.P., Lee, G., Friedman, D.A., Brachmann, E., et al. (2018) Foraging behavior and locomotion of the invasive Argentine ant from winter aggregations. PLOS ONE. 13 (8), e0202117. 10.1371/journal.pone.0202117.

15. Calderai, G., Baggiani, B., Bianchi, S., Pierucci, V., et al. (2023) Nectar-borne GABA promotes flower fidelity in bumble bees. Entomologia Generalis. 10.1127/entomologia/2023/2062.

16. Cammaerts, M., Rachidi, Z. & Gosset, G. (2014) Physiological and ethological effects of caffeine, theophylline, cocaine and atropine; Study using the ant *Myrmica sabuleti* (Hymenoptera, Formicidae) as a biological model. International Journal of Biology. 6 (3). 10.5539/ijb.v6n3p64.

17. Carlesso, D., Smargiassi, S., Pasquini, E., Bertelli, G., et al. (2021) Nectar non-protein amino acids (NPAAs) do not change nectar palatability but enhance learning and memory in honey bees. Scientific Reports. 11 (1). 10.1038/s41598-021-90895-z.

18. Cartwright, B.A. & Collett, T.S. (1987) Landmark maps for honeybees. Biological Cybernetics. 57 (1–2), 85–93. 10.1007/BF00318718.

19. Collett, M., Chittka, L. & Collett, T.S. (2013) Spatial memory in insect navigation. Current Biology. 23 (17), 789–800. 10.1016/j.cub.2013.07.020.

20. Collett, T.S. (1996) Insect navigation en route to the goal: Multiple strategies for the use of landmarks. Journal of Experimental Biology. 199 (1), 227–235. 10.1242/jeb.199.1.227.

21. Collett, T.S. & Cartwright, B.A. (1983) Eidetic images in insects: Their role in navigation. Trends in Neurosciences. 6 (3), 101–105. https://psycnet.apa.org/doi/10.1016/0166-2236(83)90048-6.

22. Collett, T.S., Dillmann, E., Giger, A. & Wehner, R. (1992) Visual landmarks and route following in desert ants. Journal of Comparative Physiology A. 170, 435–442. 10.1007/BF00191460.

23. Collett, T.S. & Zeil, J. (2018) Insect learning flights and walks. Current Biology. 28 (17), 984– 988. 10.1016/j.cub.2018.04.050.

24. Couvillon, M.J., Toufailia, H.A., Butterfield, T.M., Schrell, F., et al. (2015) Caffeinated forage tricks honeybees into increasing foraging and recruitment behaviors. Current Biology. 25 (21), 2815–2818. 10.1016/j.cub.2015.08.052.

25. Czaczkes, T.J., Grüter, C. & Ratnieks, F.L.W. (2013) Negative feedback in ants: crowding results in less trail pheromone deposition. Journal of The Royal Society Interface. 10 (81). 10.1098/rsif.2012.1009.

26. Czaczkes, T.J., Schlosser, L., Heinze, J. & Witte, V. (2014) Ants use directionless odour cues to recall odour-associated locations. Behavioral Ecology and Sociobiology. 68 (6), 981–988. 10.1007/s00265-014-1710-2.

27. De Agrò, M., Oberhauser, F.B., Loconsole, M., Galli, G., et al. (2020) Multi-modal cue integration in the black garden ant. Animal Cognition. 23 (6), 1119–1127. 10.1007/s10071-020-01360-9.

28. Deeti, S. & Cheng, K. (2021) Learning walks in an Australian desert ant, *Melophorus bagoti*. Journal of Experimental Biology. 224 (16), jeb242177. 10.1242/jeb.242177.

29. Detzel, A. & Wink, M. (1993) Attraction, deterrence or intoxication of bees (*Apis mellifera*) by plant allelochemicals. Chemoecology. 4 (1), 8–18. 10.1007/BF01245891.

30. Felsenberg, J., Gehring, K.B., Antemann, V. & Eisenhardt, D. (2011) Behavioural pharmacology in classical conditioning of the proboscis extension response in honeybees (Apis mellifera). Journal of Visualized Experiments. (47), 2282. 10.3791/2282.

31. Fewell, J.H. (1988) Energetic and time costs of foraging in harvester ants, *Pogonomyrmex occidentalis*. Behavioral Ecology and Sociobiology. 22, 401–408. 10.1007/BF00294977.

32. Fleischmann, P.N., Christian, M., Müller, V.L., Rössler, W., et al. (2016) Ontogeny of learning walks and the acquisition of landmark information in desert ants, *Cataglyphis fortis*. Journal of Experimental Biology. 219 (19), 3137–3145. 10.1242/jeb.140459.

33. Fox, J. & Weisberg, S. (2019) An R companion to applied regression. Third.. Thousand Oaks CA, Sage. Available from: https://socialsciences.mcmaster.ca/jfox/Books/Companion/.

34. Galante, H. & Czaczkes, T.J. (2023) Invasive ant learning is not affected by seven potential neuroactive chemicals. Current Zoology. 10.1093/cz/zoad001.

35. Gold, P.E. (2003) Acetylcholine modulation of neural systems involved in learning and memory. Neurobiology of Learning and Memory. 80 (3), 194–210. 10.1016/j.nlm.2003.07.003.

36. Hartig, F. (2022) DHARMa: Residual diagnostics for hierarchical (multi-level/mixed) regression models. Available from: https://CRAN.R-project.org/package=DHARMa.

37. Haubrock, P.J., Turbelin, A.J., Cuthbert, R.N., Novoa, A., et al. (2021) Economic costs of invasive alien species across Europe. NeoBiota. 67, 153–190. 10.3897/neobiota.67.58196.

38. Heinze, S., Narendra, A. & Cheung, A. (2018) Principles of insect path integration. Current Biology. 28 (17), 1043–1058. 10.1016/j.cub.2018.04.058.

39. Helmy, O. & Jander, R. (2003) Topochemical learning in black carpenter ants (*Camponotus pennsylvanicus*). Insectes Sociaux. 50 (1), 32–37. 10.1007/s000400300005.

40. Hoffmann, B.D., Luque, G.M., Bellard, C., Holmes, N.D., et al. (2016) Improving invasive ant eradication as a conservation tool: A review. Biological Conservation. 198, 37–49. 10.1016/j.biocon.2016.03.036.

41. Hogg, B.N., Nelson, E.H., Hagler, J.R. & Daane, K.M. (2018) Foraging distance of the Argentine ant in California vineyards. Journal of Economic Entomology. 111 (2), 672–679. 10.1093/jee/tox366.

42. Holway, D.A., Lach, L., Suarez, A.V., Tsutsui, N.D., et al. (2002) The causes and consequences of ant invasions. Annual Review of Ecology and Systematics. 33 (1), 181–233. 10.1146/annurev.ecolsys.33.010802.150444.

43. Huber, R. & Knaden, M. (2018) Desert ants possess distinct memories for food and nest odors. Proceedings of the National Academy of Sciences. 115 (41), 10470–10474. 10.1073/pnas.1809433115.

44. Huber, R. & Knaden, M. (2017) Homing ants get confused when nest cues are also route cues. Current Biology. 27, 3706–3710. 10.1016/j.cub.2017.10.039.

45. Humphrey, B., Helton, W.S. & Nelson, X.J. (2019) Caffeine affects the vigilance decrement of *Trite planiceps* jumping spiders (Salticidae). Journal of Comparative Psychology. 133 (4), 551–557. 10.1037/com0000203.

46. Josens, R., Eschbach, C. & Giurfa, M. (2009) Differential conditioning and long-term olfactory memory in individual *Camponotus fellah* ants. Journal of Experimental Biology. 212 (12), 1904–1911. 10.1242/jeb.030080.

47. Kaplan, G.B., Greenblatt, D.J., Ehrenberg, B.L., Goddard, J.E., et al. (1997) Dose-dependent pharmacokinetics and psychomotor effects of caffeine in humans. The Journal of Clinical Pharmacology. 37 (8), 693–703. 10.1002/j.1552-4604.1997.tb04356.x.

48. Kassambara, A., Kosinski, M. & Biecek, P. (2021) *survminer: Drawing survival curves using ‘ggplot2’*.. Available from: https://CRAN.R-project.org/package=survminer.

49. Knaden, M. & Graham, P. (2016) The sensory ecology of ant navigation: From natural environments to neural mechanisms. Annual Review of Entomology. 61 (1), 63–76. 10.1146/annurev-ento-010715-023703.

50. Lach, L., Volp, T.M. & Wilder, S.M. (2019) Previous diet affects the amount but not the type of bait consumed by an invasive ant. Pest Management Science. 75 (10), 2627–2633. 10.1002/ps.5365.

51. Lenth, R.V. (2022) *emmeans: Estimated marginal means, aka least-squares means*.. Available from: https://CRAN.R-project.org/package=emmeans.

52. Lowe, S., Browne, M., Boudjelas, S. & Poorter, M.D. (2000) 100 of the world’s worst invasive alien species: A selection from the global invasive species database. The Invasive Species Specialist Group. 12 (1). Available from: https://portals.iucn.org/library/sites/library/files/documents/2000-126.pdf.

53. Madsen, N.E.L. & Offenberg, J. (2019) Effect of caffeine and γ-aminobutyric acid on preference for sugar solutions in two ant species. Asian Myrmecology. 11. 10.20362/am.011003.

54. Majid, A.H.A., Dieng, H., Ellias, S.E., Sabtu, F.S., et al. (2018) Olfactory behavior and response of household ants (Hymenoptera) to different types of coffee odor: A coffee-based bait development prospect. Journal of Asia-Pacific Entomology. 21 (1), 46–51. 10.1016/j.aspen.2017.11.005.

55. Mathis, A., Mamidanna, P., Cury, K.M., Abe, T., et al. (2018) DeepLabCut: Markerless pose estimation of user-defined body parts with deep learning. Nature Neuroscience. 21, 1281– 1289. 10.1038/s41593-018-0209-y.

56. Menchetti, M., Schifani, E., Alicata, A., Cardador, L., et al. (2023) The invasive ant *Solenopsis invicta* is established in Europe. Current Biology. 33 (17), 896–897. 10.1016/j.cub.2023.07.036.

57. Merrill, K.C., Boser, C.L., Hanna, C., Holway, D.A., et al. (2018) Argentine ant (*Linepithema humile*, Mayr) eradication efforts on San Clemente Island, California, USA. Western North American Naturalist. 78 (4), 829–836. 10.3398/064.078.0422.

58. Müller, M. & Wehner, R. (2010) Path integration provides a scaffold for landmark learning in desert ants. Current Biology. 20 (15), 1368–1371. 10.1016/j.cub.2010.06.035.

59. Müller, M. & Wehner, R. (1994) The hidden spiral: Systematic search and path integration in desert ants, Cataglyphis fortis. Journal of Comparative Physiology A. 175, 525–530. 10.1007/BF00199474.

60. Müller, M. & Wehner, R. (2007) Wind and sky as compass cues in desert ant navigation. Naturwissenschaften. 94 (7), 589–594. 10.1007/s00114-007-0232-4.

61. Mustard, J.A. (2020) Neuroactive nectar: Compounds in nectar that interact with neurons. Arthropod-Plant Interactions. 14 (2), 151–159. 10.1007/s11829-020-09743-y.

62. Mustard, J.A. (2014) The buzz on caffeine in invertebrates: Effects on behavior and molecular mechanisms. Cellular and Molecular Life Sciences. 71 (8), 1375–1382. 10.1007/s00018-013-1497-8.

63. Mustard, J.A., Dews, L., Brugato, A., Dey, K., et al. (2012) Consumption of an acute dose of caffeine reduces acquisition but not memory in the honey bee. Behavioural Brain Research. 232 (1), 217–224. 10.1016/j.bbr.2012.04.014.

64. Muth, F., Philbin, C.S., Jeffrey, C.S. & Leonard, A.S. (2022) Discovery of octopamine and tyramine in nectar and their effects on bumblebee behavior. iScience. 25 (8). 10.1016/j.isci.2022.104765.

65. Narendra, A., Reid, S.F. & Hemmi, J.M. (2010) The twilight zone: Ambient light levels trigger activity in primitive ants. Proceedings of the Royal Society B: Biological Sciences. 277 (1687), 1531–1538. 10.1098/rspb.2009.2324.

66. Nath, T., Mathis, A., Chen, A.C., Patel, A., et al. (2019) Using DeepLabCut for 3D markerless pose estimation across species and behaviors. Nature Protocols. 14, 2152–2176. 10.1038/s41596-019-0176-0.

67. Nicolson, S.W. (2022) Sweet solutions: nectar chemistry and quality. Philosophical Transactions of the Royal Society B: Biological Sciences. 377 (1853). 10.1098/rstb.2021.0163.

68. Nyamukondiwa, C. & Addison, P. (2011) Preference of foraging ants (Hymenoptera: Formicidae) for bait toxicants in South African vineyards. Crop Protection. 30 (8), 1034–1038. 10.1016/j.cropro.2011.03.014.

69. Oberhauser, F.B., Schlemm, A., Wendt, S. & Czaczkes, T.J. (2019) Private information conflict: *Lasius niger* ants prefer olfactory cues to route memory. Animal Cognition. 22 (3), 355–364. 10.1007/s10071-019-01248-3.

70. Pallarés, J.G., Fernández-Elías, V.E., Ortega, J.F., Muñoz, G., et al. (2013) Neuromuscular responses to incremental caffeine doses: Performance and side effects. Medicine & Science in Sports & Exercise. 45 (11), 2184–2192. 10.1249/MSS.0b013e31829a6672.

71. Piqueret, B., Bourachot, B., Leroy, C., Devienne, P., et al. (2022) Ants detect cancer cells through volatile organic compounds. iScience. 25 (3), 103959. 10.1016/j.isci.2022.103959.

72. Popp, S. & Dornhaus, A. (2023) Ants combine systematic meandering and correlated random walks when searching for unknown resources. iScience. 26 (2), 105916. 10.1016/j.isci.2022.105916.

73. R Core Team (2022) R: A language and environment for statistical computing.. Vienna, Austria, R Foundation for Statistical Computing. Available from: https://www.R-project.org/.

74. Reid, C.R., Latty, T. & Beekman, M. (2012) Making a trail: informed Argentine ants lead colony to the best food by U-turning coupled with enhanced pheromone laying. Animal Behaviour. 84 (6), 1579–1587. 10.1016/j.anbehav.2012.09.036.

75. Reid, C.R., Sumpter, D.J.T. & Beekman, M. (2011) Optimisation in a natural system: Argentine ants solve the Towers of Hanoi. Journal of Experimental Biology. 214 (1), 50–58. 10.1242/jeb.048173.

76. Roces, F. (1990) Olfactory conditioning during the recruitment process in a leaf-cutting ant. Oecologia. 83 (2), 261–262. 10.1007/BF00317762.

77. Rossi, N., Pereyra, M., Moauro, M.A., Giurfa, M., et al. (2020) Trail-pheromone modulates subjective reward evaluation in Argentine ants. Journal of Experimental Biology. 223 (17). 10.1242/jeb.230532.

78. Rössler, W. (2023) Multisensory navigation and neuronal plasticity in desert ants. Trends in Neurosciences. 46 (6), 415–417. 10.1016/j.tins.2023.03.008.

79. Rossum, G.V. & Drake, F.L. (2009) Python3 reference manual. Scotts Valley, CA, CreateSpace.

80. Rust, M.K., Reierson, D.A. & Klotz, J.H. (2003) Pest management of Argentine ants (Hymenoptera: Formicidae). Journal of Entomological Science. 38 (2), 159–169. 10.18474/0749-8004-38.2.159.

81. Scheiner, R., Plückhahn, S., Öney, B., Blenau, W., et al. (2002) Behavioural pharmacology of octopamine, tyramine and dopamine in honey bees. Behavioural Brain Research. 136 (2), 545–553. 10.1016/S0166-4328(02)00205-X.

82. Schilman, P.E., Lighton, J.R.B. & Holway, D.A. (2005) Respiratory and cuticular water loss in insects with continuous gas exchange: Comparison across five ant species. Journal of Insect Physiology. 51 (12), 1295–1305. 10.1016/j.jinsphys.2005.07.008.

83. Schultheiss, P., Wystrach, A., Legge, E.L.G. & Cheng, K. (2012) Information content of visual scenes influences systematic search of desert ants. Journal of Experimental Biology. 216, 742– 749. 10.1242/jeb.075077.

84. Schwarz, S., Wystrach, A. & Cheng, K. (2017) Ants’ navigation in an unfamiliar environment is influenced by their experience of a familiar route. Scientific Reports. 7 (1), 14161. 10.1038/s41598-017-14036-1.

85. Si, A., Zhang, S. & Maleszka, R. (2005) Effects of caffeine on olfactory and visual learning in the honey bee (*Apis mellifera*). Pharmacology Biochemistry and Behavior. 82 (4), 664–672. 10.1016/j.pbb.2005.11.009.

86. Silverman, J. & Brightwell, R.J. (2008) The Argentine ant: Challenges in managing an invasive unicolonial pest. Annual Review of Entomology. 53 (1), 231–252. 10.1146/annurev.ento.53.103106.093450.

87. Singaravelan, N., Nee’man, G., Inbar, M. & Izhaki, I. (2005) Feeding responses of free-flying honeybees to secondary compounds mimicking floral nectars. Journal of Chemical Ecology. 31 (12), 2791–2804. 10.1007/s10886-005-8394-z.

88. Smith, A. (2002) Effects of caffeine on human behavior. Food and Chemical Toxicology. 40 (9), 1243–1255. 10.1016/S0278-6915(02)00096-0.

89. Souza, J.G., Coso, J., Fonseca, F.S., Silva, B.V.C., et al. (2022) Risk or benefit? Side effects of caffeine supplementation in sport: A systematic review. European Journal of Nutrition. 61 (8), 3823–3834. 10.1007/s00394-022-02874-3.

90. Steck, K., Hansson, B.S. & Knaden, M. (2011) Desert ants benefit from combining visual and olfactory landmarks. Journal of Experimental Biology. 214 (8), 1307–1312. 10.1242/jeb.053579.

91. Steck, K., Hansson, B.S. & Knaden, M. (2009) Smells like home: Desert ants, *Cataglyphis fortis*, use olfactory landmarks to pinpoint the nest. Frontiers in Zoology. 6 (1), 5. 10.1186/1742-9994-6-5.

92. Suiter, D.R., Gochnour, B.M., Holloway, J.B. & Vail, K.M. (2021) Alternative methods of ant (Hymenoptera: Formicidae) control with emphasis on the Argentine ant, *Linepithema humile*. Insects. 12 (6), 487. 10.3390/insects12060487.

93. Sumpter, D.J.T. & Beekman, M. (2003) From nonlinearity to optimality: pheromone trail foraging by ants. Animal Behaviour. 66 (2), 273–280. 10.1006/anbe.2003.2224.

94. Therneau, T.M. (2022) coxme: Mixed effects cox models.. Available from: https://CRAN.R-project.org/package=coxme.

95. Therneau, T.M. & Grambsch, P.M. (2000) *Modeling survival data: Extending the cox model*. New York, Springer.

96. Thomson, J.D., Draguleasa, M.A. & Tan, M.G. (2015) Flowers with caffeinated nectar receive more pollination. Arthropod-Plant Interactions. 9 (1), 1–7. 10.1007/s11829-014-9350-z.

97. Tiedeken, E.J., Stout, J.C., Stevenson, P.C. & Wright, G.A. (2014) Bumblebees are not deterred by ecologically relevant concentrations of nectar toxins. Journal of Experimental Biology. 217 (9). 10.1242/jeb.097543.

98. Van Swinderen, B. & Andretic, R. (2011) Dopamine in *Drosophila*: setting arousal thresholds in a miniature brain. Proceedings of the Royal Society B: Biological Sciences. 278 (1707), 906– 913. 10.1098/rspb.2010.2564.

99. Vega, S.Y. & Rust, M.K. (2003) Determining the foraging range and origin of resurgence after treatment of Argentine ant (Hymenoptera: Formicidae) in urban areas. Journal of Economic Entomology. 96 (3), 844–849. 10.1093/jee/96.3.844.

100. Von Thienen, W., Metzler, D., Choe, D. & Witte, V. (2014) Pheromone communication in ants: a detailed analysis of concentration-dependent decisions in three species. Behavioral Ecology and Sociobiology. 68 (10), 1611–1627. 10.1007/s00265-014-1770-3.

101. Wagner, T., Galante, H., Josens, R. & Czaczkes, T.J. (2023) Systematic examination of learning in the invasive ant *Linepithema humile* reveals fast learning and long-lasting memory. Animal Behaviour. 203, 41–52. 10.1016/j.anbehav.2023.06.012.

102. Wajnberg, E., Acosta-Avalos, D., Alves, O.C., Oliveira, J.F., et al. (2010) Magnetoreception in eusocial insects: An update. Journal of The Royal Society Interface. 7 (2), 207–225. 10.1098/rsif.2009.0526.focus.

103. Wehner, R. (2008) The desert ant’s navigational toolkit: Procedural rather than positional knowledge. Navigation. 55 (2), 101–114. 10.1002/j.2161-4296.2008.tb00421.x.

104. Wehner, R., Marsh, A.C. & Wehner, S. (1992) Desert ants on a thermal tightrope. Nature. 357 (6379), 586–587. 10.1038/357586a0.

105. Wehner, R., Meier, C. & Zollikofer, C. (2004) The ontogeny of foraging behaviour in desert ants, *Cataglyphis bicolor*. Ecological Entomology. 29 (2), 240–250. 10.1111/j.0307-6946.2004.00591.x.

106. Wehner, R. & Müller, M. (2006) The significance of direct sunlight and polarized skylight in the ant’s celestial system of navigation. Proceedings of the National Academy of Sciences. 103 (33), 12575–12579. 10.1073/pnas.0604430103.

107. Wickham, H. (2016) ggplot2: Elegant graphics for data analysis. Springer-Verlag New York. Available from: https://ggplot2.tidyverse.org.

108. Wickham, H. (2022) *stringr: Simple, consistent wrappers for common string operations*.. R package version 1.5.0. Available from: https://CRAN.R-project.org/package=stringr.

109. Wink, M. (2018) Plant secondary metabolites modulate insect behavior-steps toward addiction? Frontiers in Physiology. 9, 364. 10.3389/fphys.2018.00364.

110. Wittlinger, M., Wehner, R. & Wolf, H. (2006) The ant odometer: Stepping on stilts and stumps. Science. 312 (5782), 1965–1967. 10.1126/science.1128659.

111. Wright, G.A., Baker, D.D., Palmer, M.J., Stabler, D., et al. (2013) Caffeine in floral nectar enhances a pollinator’s memory of reward. Science. 339 (6124), 1202–1204. 10.1126/science.1228806.

112. Wystrach, A. & Beugnon, G. (2009) Ants learn geometry and features. Current Biology. 19 (1), 61–66. 10.1016/j.cub.2008.11.054.

113. Wystrach, A., Beugnon, G. & Cheng, K. (2011) Landmarks or panoramas: What do navigating ants attend to for guidance? Frontiers in Zoology. 8 (1), 21. 10.1186/1742-9994-8-21.

114. Wystrach, A., Buehlmann, C., Schwarz, S., Cheng, K., et al. (2020) Rapid aversive and memory trace learning during route navigation in desert ants. Current Biology. 30 (10), 1927–1933. 10.1016/j.cub.2020.02.082.

115. Wystrach, A., Schwarz, S., Schultheiss, P., Baniel, A., et al. (2014) Multiple sources of celestial compass information in the Central Australian desert ant *Melophorus bagoti*. Journal of Comparative Physiology A. 200 (6), 591–601. 10.1007/s00359-014-0899-x.

116. Yates, A.A. & Nonacs, P. (2016) Preference for straight-line paths in recruitment trail formation of the Argentine ant, *Linepithema humile*. Insectes Sociaux. 63 (4), 501–505. 10.1007/s00040-016-0492-0.

117. Yeoh, X.L., Dieng, H. & Majid, A.H.A. (2018) Mortality and repellent effects of coffee extracts on the workers of three household ant species. Pertanika Journal of Tropical Agricultural Science. 41 (4), 1557–1586.

118. Zeil, J. (2022) Visual navigation: Properties, acquisition and use of views. Journal of Comparative Physiology A. 10.1007/s00359-022-01599-2.

119. Zeil, J. & Fleischmann, P.N. (2019) The learning walks of ants (Hymenoptera: Formicidae). Myrmecological News. 29, 93–110. 10.25849/MYRMECOL.NEWS_029:093.

120. Zeil, J., Hofmann, M.I. & Chahl, J.S. (2003) Catchment areas of panoramic snapshots in outdoor scenes. Journal of the Optical Society of America A. 20 (3), 450. 10.1364/JOSAA.20.000450.

121. Zeil, J., Narendra, A. & Stürzl, W. (2014) Looking and homing: how displaced ants decide where to go. Philosophical Transactions of the Royal Society B: Biological Sciences. 369 (1636). 10.1098/rstb.2013.0034.

